# A multiscale modeling framework for transport of PEGylated lipid nanoparticle through the extracellular matrix

**DOI:** 10.64898/2026.09.09.750406

**Authors:** Prasheel Nakate, Kyle J. Colston, Severin T. Schneebeli, Arezoo M. Ardekani

## Abstract

Lipid nanoparticles (LNPs) are one of the leading platforms for delivering nucleic acid therapeutics, yet their efficacy is limited by physicochemical interactions with the extracellular matrix (ECM) that trap the particles before they reach target cells. PEGylated nanoparticles mitigate these interactions by forming a protective steric layer on their surfaces. However, there is a lack of a predictive tool that gives mechanistic insights about how PEG surface density governs the underlying interaction and results in enhanced diffusive transport of LNPs through the ECM. Here, we present a multi-scale hierarchical computational framework that couples all-atom constant pH molecular dynamics (CpHMD) with a highly coarse-grained model of the complete LNP within a crosslinked hyaluronic acid (HA) network. These atomistic simulations resolve the free energy of interaction between the LNP surface and HA chains across varying PEG lipid compositions, and integrate these free energy profiles to inform the coarse-grained simulations of LNP transport through the matrix. This work highlights that even a slightly PEGylated surface depletes the near-contact shell between the LNP and HA chains, which disrupts their adhesive interactions. These protective PEG layers produce a sharp, non-linear enhancement in LNP diffusivities, with just 1% PEG increasing the diffusivity nearly eight-fold relative to bare LNPs, which remain trapped in the matrix structures. This work provides a quantitative estimate of how PEG surface density governs LNP transport through the ECM, which offers predictive guidance for engineering LNP surface properties in target-specific drug delivery.

## Introduction

Over the past couple of decades, nanoparticle-based therapeutics have emerged as a promising platform for delivering drugs to complex target sites. Specifically, lipid-based nanoparticles (LNPs) offer great capability to deliver nucleic acid-based therapeutics. These LNPs not only protect the cargo present in their core, but also enable efficient release at targeted cells ^1–3^. Despite their therapeutic advantage, these LNPs are susceptible to clearance by the phagocytic system, which recognizes them as foreign particles ^4,5^. Such an event results in poor circulation, half-life, and rapid elimination of LNPs from the body ^6,7^. Additionally, bare LNPs interact with multiple binding sites during their transport in the extracellular matrix (ECM) before reaching the target cells. These surface interactions with the extracellular matrix result in a significant fraction of LNPs becoming trapped within the matrix structure ^8,9^.

The behavior of nanoparticles in biological systems is heavily influenced by their surface interactions. In order to improve the circulation half-life of these nanoparticles, they are functionalized with polyethylene glycol (PEG) ^10,11^. These PEG molecules form a stealth coating on the surface of nanoparticles that significantly restricts the attractive interactions of the nanoparticle surface with other entities in the biological system ^12,13^. A lipid nanoparticle is typically composed of ionizable lipid, helper lipid, cholesterol, and PEGylated lipid (PEG-lipid) ^14^. Each of these components serves a critical role in the delivery of the LNPs to the target sites. Generally, the composition of LNPs involves around 50% ionizable lipid, 10% helper lipid, 38.5% cholesterol, and 1.5%PEG-lipids ^15^. Among these components, PEGylated lipids are incorporated into the LNP membrane such that their hydrophobic lipid moieties associate with other lipids in the membrane, while the hydrophilic PEG chains extend outward, forming a hydration layer. This protective layer reduces clearance from the phagocytic system, binding with matrix structures, and aggregation in the solution state through steric interactions ^16,17^.

There are several critical factors that dictate the performance of stealth-coated nanoparticles in biological systems. These include the molecular weight, surface density, and chain conformations of PEGylated lipid molecules ^18–20^. As the molecular weight of the PEG increases, their chain length and the thickness of the hydration layer proportionately increase. In a study conducted by Gref *et al*., it was shown that increasing PEG molecular weight from 2 to 5 kDa resulted in a significant reduction in the clearance of nanoparticles. Beyond 5 kDa molecular weight, the effect of PEG chain length is not significant ^21^ as surface density and chain conformation start influencing the structure of the protective layer. In general, it is established that a minimum of 2 kDa PEG molecular weight is required to create a stable protective layer with controlled flexibility of the chains ^7^. For lipid nanoparticles, the chain length of the lipid moieties of PEG-lipids also plays a crucial role ^22^. The 14-carbon (C-14) chain PEG-lipids are loosely bound in the lipid membrane and prone to quick shedding from the LNPs ^23^. This desorption results in lower surface coverage of PEG chains and poor circulation half-life of LNPs. Whereas for the 18-carbon (C-18) chain PEG-lipids, more energy is required to remove them from the membrane due to the stronger hydrophobic interactions. Therefore, the C-18 LNPs show better circulation half-life due to slower shedding from the surface ^23^.

The physical characteristics of this steric layer such as thickness and continuity influences the transport of LNPs in the matrix structure. If the layer is patchy and sparse, it partially exposes the LNP surface to the HA polymer chains. Whereas a continuous thick layer strongly repels the polymeric structures assisting transport of LNPs. The characteristics of PEG chains such as surface density and their chain conformations are interlinked with each other ^24^. The distance between the PEG grafts *D*, and the Flory radius (coil size) *R*_*F*_ of the chain dictate the dominant chain conformation on the nanoparticle surface ^25^. At lower surface densities, the PEG chains are well separated where *D > R*_*F*_ , and they are not fully extended from the particle surface, which creates a “mushroom” conformation ^26^. On the other hand, when surface density is higher (*D < R*_*F*_ ), the chains start to overlap, pushing them to extend outwards from the nanoparticle surface. In this case, PEG chains form a “brush” configuration, resulting in a thicker hydration layer on the nanoparticle surface ^27^.

One of the key hurdles in the delivery of nanoparticle-based therapeutics is transport through the extracellular matrix. The extent of matrix penetration is strongly influenced by nanocarrier design parameters. To date, interactions between extracellular matrix components and LNP surfaces have been characterized primarily through empirical correlations, often linked to PEG surface density. However, a predictive, mechanistic understanding, particularly in terms of the underlying free energy landscape governing these interactions and its overall impact on the diffusive transport of LNPs, remains largely unclear. In this work, we present a multiscale hierarchical computational framework to accurately resolve the interactions between the hyaluronic acid (HA) polymer chains in the ECM and the PEGylated LNP surface. Here, we perform all-atom constant pH molecular dynamics (CpHMD) simulations for the LNP surfaces involving different PEG surface densities to capture the free energy between the LNP surface and the hyaluronic acid chains. We feed these energy profiles to our previously developed highly coarse-grained molecular dynamics computational framework that models the whole LNP structure inside the ECM matrix environment ^28^. With this multiscale modeling framework, we quantify the effect of variable PEG densities on the transport of LNPs in the extracellular matrix structures. This study aims at providing crucial predictive insights for engineering LNP surface properties for target-specific drug delivery.

## Methods

To investigate the effect of different PEG-lipid compositions in the LNP membranes on overall transport through the ECM structures, we perform multiscale hierarchical molecular simulations. Here, we first calculate the highly accurate free energy profile between the two key interacting components in the transport processes. This involves the hyaluronic acid polymer chain present abundantly in the ECM structures and the surface of the LNP membrane coated with pegylated lipids. To model these lipid membranes, we consider DLin-DMA-MC3 (ionizable lipid), cholesterol, DSPC (helper lipid), and DMG PEG-2000 (PEG lipid) molecules. To perform this study, we consider three representative cases of PEG-lipid compositions: 0%, 1%, and 5% as shown in Figure 1.

**Fig. 1.**
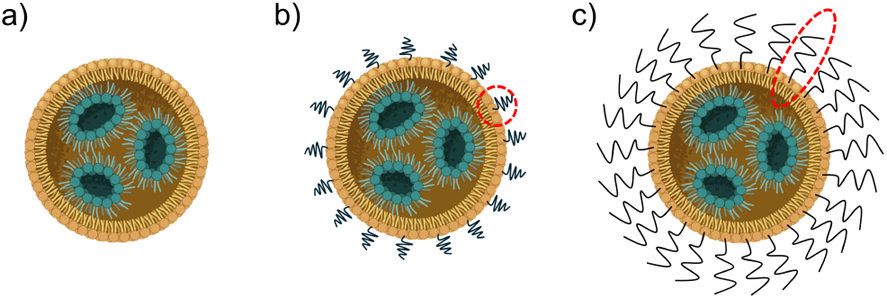
Schematic of LNP structures with different PEG composition in the membrane: a) Bare LNP with 0% PEG-lipids, b) LNP with 1% PEG-lipids in “mushroom” conformation, and c) LNP with 5% PEG-lipids in “brush-like” conformation due to steric interactions.

We perform all-atom simulations using a modified version of *GROMACS 2021* that incorporates a scalable constant *p*H (CpHMD) methodology ^29^ implemented with *phbuilder* ^30^. CpHMD methods have been implemented to accurately model dynamic protonation events of ionizable sites in molecular dynamics simulations ^31 32 33 34^. We have shown that scalable CpHMD methods can accurately capture environment-dependent *p*Ka transitions in relevant membrane models containing hundreds of ionizable groups ^35 36^. The apparent *p*Ka of an LNP formulation is crucial in determining *p*H-dependent responses ^37 38^, and must be modeled carefully. Here, we use validated CpHMD parameters for DLin-MC3-DMA as previously reported ^36^, which demonstrates *p*H-dependent structures and apparent *p*Ka’s consistent with other all-atom ^39 40 41^ and coarse grain ^42 43^ models. All components are modeled with the CHARMM36 forcefield ^44^ in addition to the water, which is modeled using the TIP3P model ^45^. Each ionizable head group is treated with *λ* -dynamics, and the dynamic protonation events are modeled by interpolation between the atomic charges of protonated and deprotonated states. All systems are minimized by an initial steepest descent minimization for 5,000 steps followed by two equilibration steps. First, *NVT* ensemble equilibration is performed using a 2 fs time step for 5,000 steps at 300K with the v-rescale temperature coupling, and then *NPT* ensemble equilibration is performed using a 2 fs time step for 5,000 steps using the c-rescale method to semiisotropically maintain a pressure of 1.0 bar with a compression constant of 4.5 *×* 10^*−*5^ nm*−*1. Short-range non-bonded interactions are held to a 1.2 nm cutoff distance with electrostatic interactions calculated using the PME method ^46^ and the constraints are imposed using the LINCS algorithm ^47^. Production simulations are performed using a 2 fs time step, and we use the same parameters used for the prior equilibration steps. The membranes are assembled using the CHARMM-GUI ^48^ membrane builder to make systems containing DLin-DMA-MC3 (50 molecules), cholesterol (40 to 38 molecules), DSPC (10 molecules), and 2K PEG lipids (0 to 5 molecules). A comprehensive list of system size and components are given in Table S1.

We simulate these model membranes for 10 ns before an “infinite” HA chain (derived from pdb code: 1HYA ^49^) and 500 buffer particles are added using the *gmx insert-molecules*. Model systems are simulated for 50 ns with position restraints placed on the HA chain. Then we perform pulling simulations using a harmonic potential to pull the center-of-mass (COM) of the HA chain to the COM of the non-PEG membrane components at a rate of pm ps^*−*1^ and a force of 1000 kJ mol^*−*1^ nm^*−*2^. Sampling windows are extracted every 0.2 nm, and umbrella sampling is performed for 50 ns at each window using a harmonic potential force of 1000 kJ mol^*−*1^ nm^*−*2^. The weighted histogram analysis method (WHAM) ^50^ is used to calculate the free energy profile along the pulling coordinate as shown in Figure2 a). Different length alkyl PEG lipids (C12, C14, and C18) are compared at a 5% PEG composition and did not demonstrate a significant difference in their interaction with the HA chain, shown Figure S1. The results shown for the different % PEG compositions in the main text are simulated using DMG-PEG2000 (C14) as the PEG lipid.

**Fig. 2.**
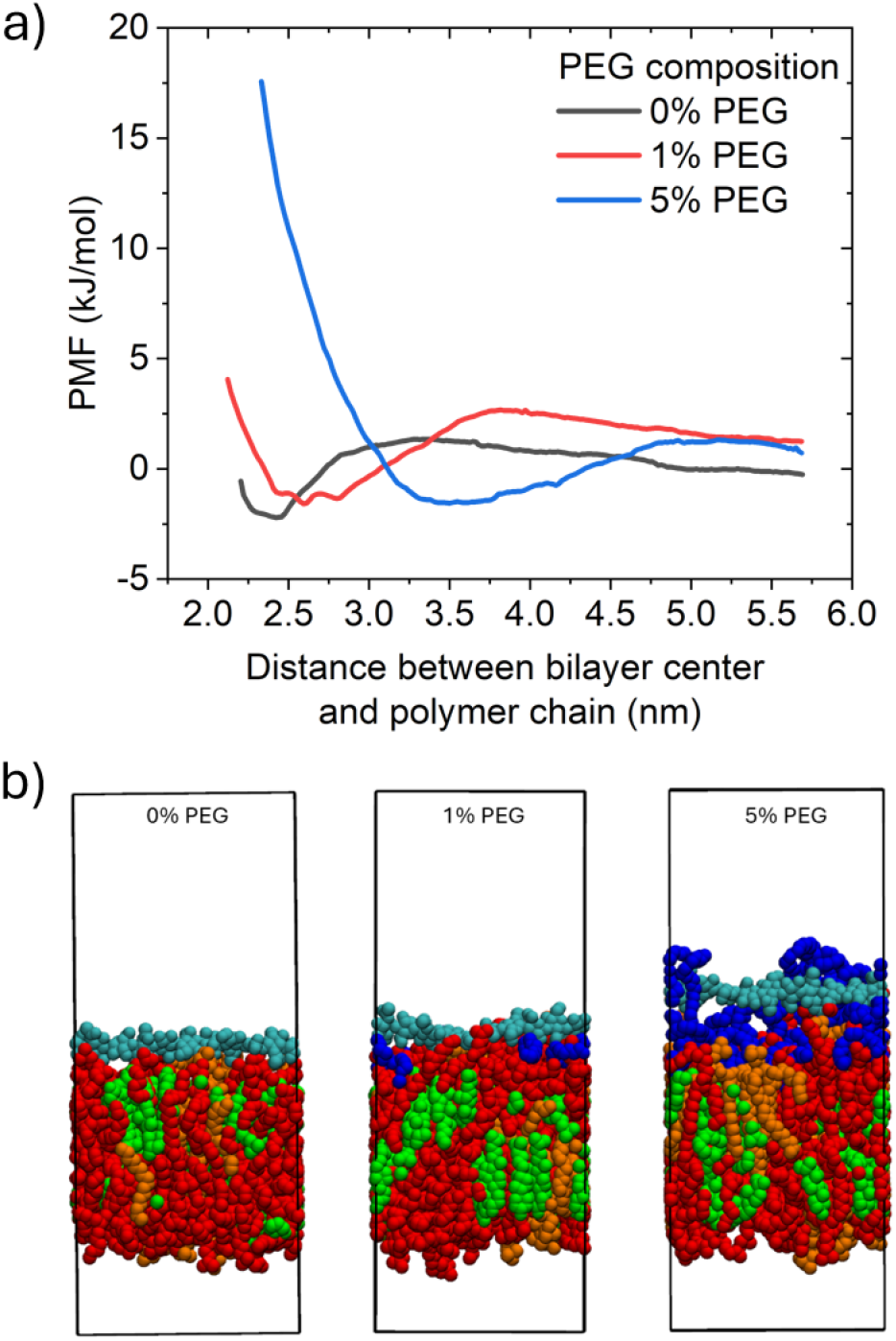
a) The free energy profiles calculated from the WHAM analysis and b) minima positions of the HA chain (cyan) relative to the center of the membrane (DLin-MC3-DMA: red, cholesterol: green, helper lipid: orange).

To evaluate the diffusive performance of the LNPs with variable PEG compositions in the matrix environment, we need the particle trajectories over longer timescales. For these calculations, we simulate the whole LNP structures in the ECM environment formed with uniformly crosslinked HA polymer chains. Our previously developed highly coarse-grained molecular simulation framework consists of LNP membrane surfaces modeled using a one-particle-thick representation and an extracellular matrix modeled with a bead-spring representation for the HA polymer chains ^28^. The one-particle-thick solvent-free representation of LNP beads significantly reduces the computational expense while preserving the key physicochemical features such as interaction with ECM structures, membrane fluidity, and topological changes ^51^. Here, the matrix structure is represented with a uniformly crosslinked polymer network with Debye interactions for electrostatic forces present due to the negatively charged HA monomeric units.

The free energy profiles in Figure 2 show strong repulsive interactions as the chain moves closer to the lipid membrane surface. At intermediate distances, there is an attraction well between these components, with the 0% PEG case showing sharper gradients in the potential well and it is closest to the bilayer center. At 0% PEG, the depth of the potential well is maximum with 2.2 kJ/mol at 2.42 nm, indicating the strongest attractive forces. As the PEG% increases, the potential well shifts away from the bi-layer center, its depth decreases slightly, and the gradients in the well become more softer, as shown in Table 1. This trend reflects a reduction in attractive interactions alongside an increase in the effective size of LNPs with increasing PEG composition. It further indicates a transition in PEG conformation from a mushroom to a brush-like regime, as illustrated in Figure 1.

**Table 1.** Features extracted from the free energy profiles between LNP surface and the HA polymer chains.

| PEG composition | Energy minimum ($\epsilon$ in kJ/mol) | Distance of HA chain from bilayer center ($r_{min}$ in nm) |
| --- | --- | --- |
| 0% | -2.20 | 2.42 |
| 1% | -1.58 | 2.60 |
| 5% | -1.56 | 3.50 |

In the present coarse-grained computational framework, the interactions between the LNP beads and the HA chain beads are modeled using a 12/6 Lennard-Jones potential shown in equation 1

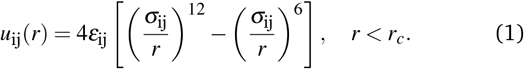

where the values of *ε*_ij_ and *σ*_ij_ are calculated using the parameters extracted from the free energy profiles shown in Table 1. As we assume the analytical form of the interaction as a 12/6 Lennard-Jones potential, we calculate 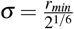, where *r*_*min*_ is the location of the energy minimum. With these calculated *σ* and *ε* values from Table 1, we map the interaction potentials between the coarse-grained LNP beads and the HA polymer chain beads as shown in Figure 3. This mapping ensures that the depth and location of the potential well in the coarse-grained description match the highly resolved all-atomistic free energy profile in Figure 2. These potentials are the key non-bonded interactions in this modeling framework. Additional details related to interaction parameter calculations are provided in the supplementary materials. The bonded interactions in the crosslinked HA polymer beads and the other non-bonded interactions between the LNP-LNP beads and the HA-HA beads are kept the same as our past study ^28^.

**Fig. 3.**
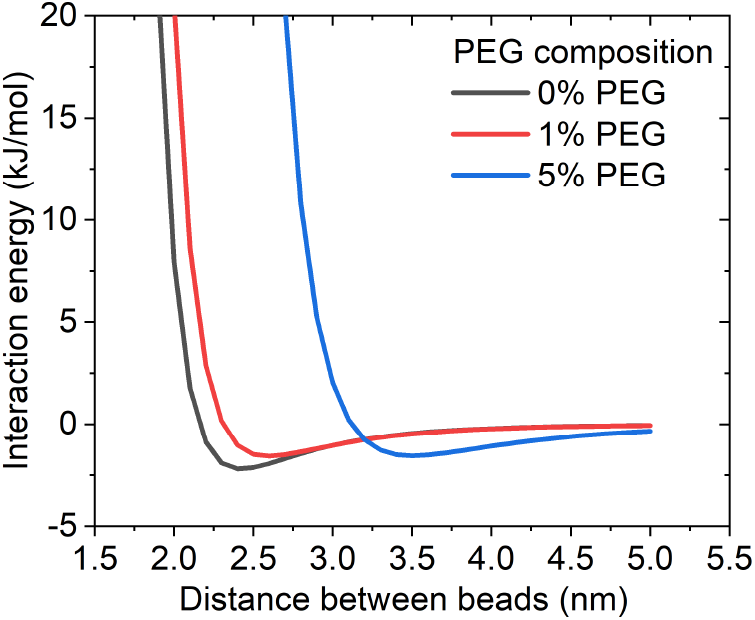
The 12/6 Lennard-Jones potential for the interaction between coarse-grained LNP beads and HA polymer chain beads

Here, we perform the coarse-grained simulations using the LAMMPS package ^52^. The system of 10 preassembled LNPs is introduced into the uniformly crosslinked ECM structure as shown in Figure 4. The equations of motion are integrated with the velocity-Verlet algorithm. We employ a Langevin thermostat with the NVE ensemble. Here, the system is kept at temperature *k*_*B*_*T* = 0.23*ε*, where *k*_*B*_ is Boltzmann’s constant. For all the cases, we perform simulations till 2 *×* 10^5^ time units, where the particle position data is stored every 100 time units. This allows us to record the trajectories for longer timescales. We further quantify the motion of particles using the mean square displacement (MSD) of the LNP centroid as it traverses the matrix structure using equation 2

**Fig. 4.**
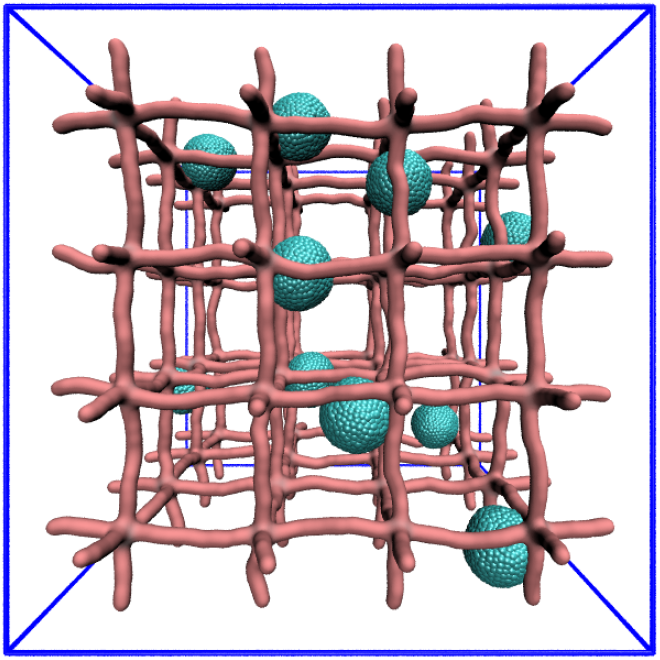
Coarse-grained computational model for the LNPs inside the uniformly crosslinked extracellular matrix structure of HA chain polymers.

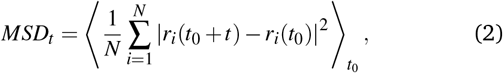

where *N* is the number of LNPs in the system, *r* represents the coordinates of the centroid of LNPs, *t* is the time lag, and *t*_0_ is the time origin.

## Results and discussion

The LNPs with 0% PEG are generally highly exposed to their surroundings. With no layer protecting their surface, they interact with the chained polymers through van der Waals forces, making them highly susceptible to sticking to the polymeric matrix. To test this, we perform the coarse-grained simulations with preassembled LNPs in the polymeric matrix structure shown in Figure 4. For all the cases of PEG compositions, we keep the distance between the crosslinking nodes uniform at 24*σ* and the size of preassembled LNP structures at 12*σ* . This keeps the particle-to-matrix mesh size ratio consistent at 50% and eliminates any size-dependent effects on the transport through the matrix structure. We simulate these LNPs in matrix with the 12/6 Lennard-Jones interaction potential given in Figure 3. We first track the particle position inside the matrix structure. Figure 5 a) shows the location of the tracer particle on the surface of a sample LNP as it tries to navigate the matrix structure. These highly concentrated trajectory lines indicate that the LNP spends most of its time near its initial position. As the tracer particle is on the surface of the LNP, the rotational motion of the LNP is also tracked, resulting in a globular nature of these trajectory lines. These lines indicate that there is almost no translational motion of these LNPs in the matrix.

**Fig. 5.**
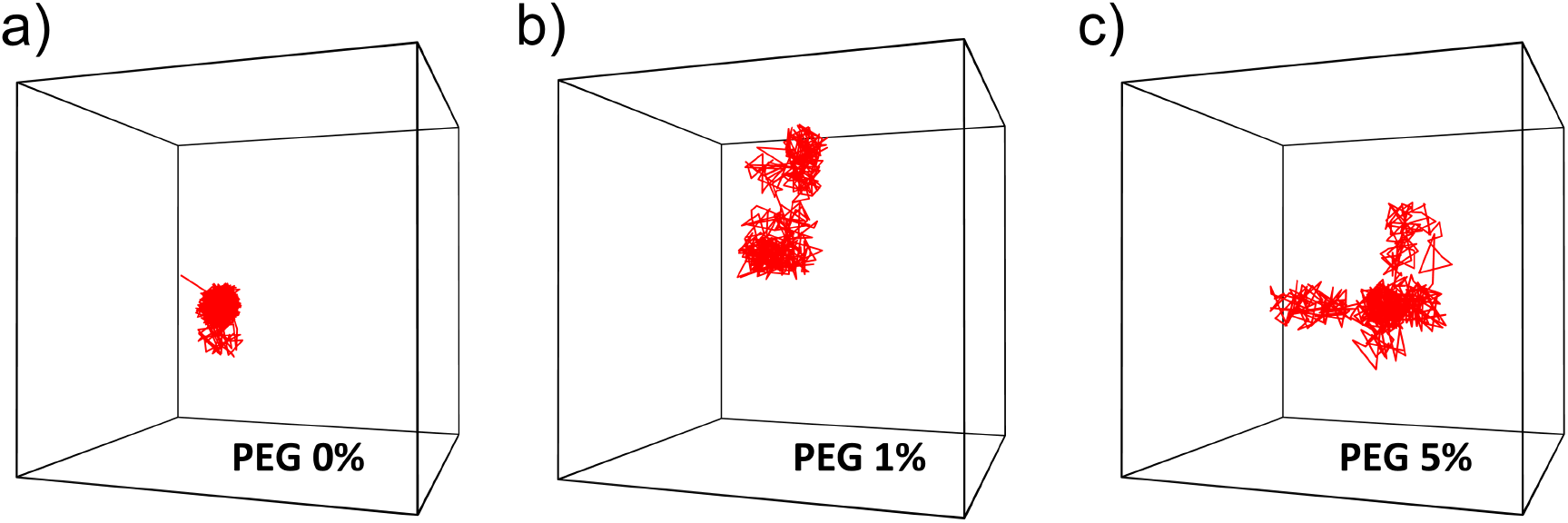
Trajectories of tracer particle on a single LNP for all the PEG compositions.

As we increase the PEG composition to 1%, there is some degree of protection layer on the surface of the LNPs. Here, the PEG chains have enough spacing among them, which creates mushroom-like structures attached to the LNP surface. In this computational framework, we model the effective surface interactions between these mushroom-coated LNPs and the polymer chain through the coarse-grained potential shown in Figure 3. Similar to the 0% PEG case, we keep the physical dimensions of the preassembled LNPs at 12*σ* . The trajectory lines shown in Figure 5 b) indicate translational motion of the LNPs in the matrix structure. As compared to the 0% PEG case, we observe widespread distribution of trajectory lines of the tracer particle. Here, the position of the LNP shifts from one caged structure to another, indicating that there is no permanent entrapment of LNPs in their initial position.

As the PEG composition approaches 5%, there is a strong layer of PEG chains surrounding the LNP structure as shown in Figure 1. This layer is so dense that it doesn’t expose the bare LNP surface to its surroundings, making it least susceptible to stick to the polymers in the matrix. The free energy profiles in Figure 2 indicate that the energy minimum is pushed away from the LNP surface at 3.5 nm. In the coarse-grained simulations, we generate the trajectory lines of these 5% PEG LNPs as shown in Figure 5 c). These lines show signs of free translational particle movement in the matrix structure. Here, the sample LNP explores multiple caged regions inside the chained network. As compared to the lines in Figure 5 a) and b), the LNPs with 5% PEG show more efficient diffusive transport through the ECM structure.

To further quantify the performance of the LNPs with different PEG compositions, we calculate their mean square displacements (MSD) as a function of time lags using equation 2. Here, we take the ensemble average of the square displacements of all ten LNPs in the system with a time lag of up to 1.8 *×* 10^5^*τ*. This ensures there is enough sample size at every time lag for accurate prediction of the mean square displacements. We perform these calculations using at least three replicate simulations to report the mean and the standard deviations in our mean square displacement predictions for all PEG compositions. Figure 6 shows the mean square displacement of LNPs with 0% PEG lipids. Here, during the initial time lags, there is a rise in the MSD, but these displacement values are related to the ballistic diffusive regime and do not indicate any long-range translational motion of the particles. The MSD curve for the 0% PEG case shows a plateauing behavior approaching 125*σ* ^2^ at higher time lags. This indicates that the long-term motion of the LNPs is arrested by the matrix. Here, the plateau height corresponds to a characteristic length of 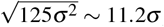 , which is roughly half of the crosslinking length 24*σ* . At the beginning of each run, LNPs are kept exactly 12*σ* distance away from the chains, which is at the center of a cage formed by surrounding polymer chains. This close agreement between the half cage width 12*σ* and the characteristic length from the plateau 11.2*σ* indicates that the dominant displacement mode of the LNPs is from the nodal center toward the surrounding polymer chain, where they become completely adsorbed.

**Fig. 6.**
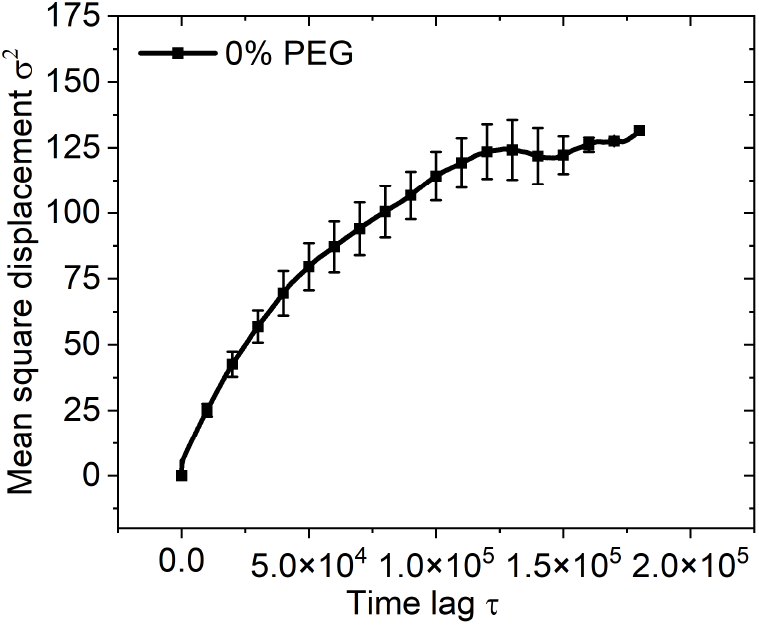
Mean square displacements of LNPs with 0% PEG as a function of time lags in the extracellular matrix.

Similarly, we compute the mean square displacements for the LNPs with 1% and 5% PEG cases as shown in Figure 7. These plots indicate that the LNPs with 1% and 5% PEG show much higher MSDs as compared to the 0% PEG case. These cases show almost linear rise in the MSDs as a function of time lags, which further indicates free translational diffusion of the LNPs in the ECM structures. The LNPs with 5% PEG reach almost 1500 *σ*^2^ of displacements towards the end of the time lags. Whereas the case with 1% PEG reaches slightly more than half of the total MSD shown by 5% PEG LNPs.

**Fig. 7.**
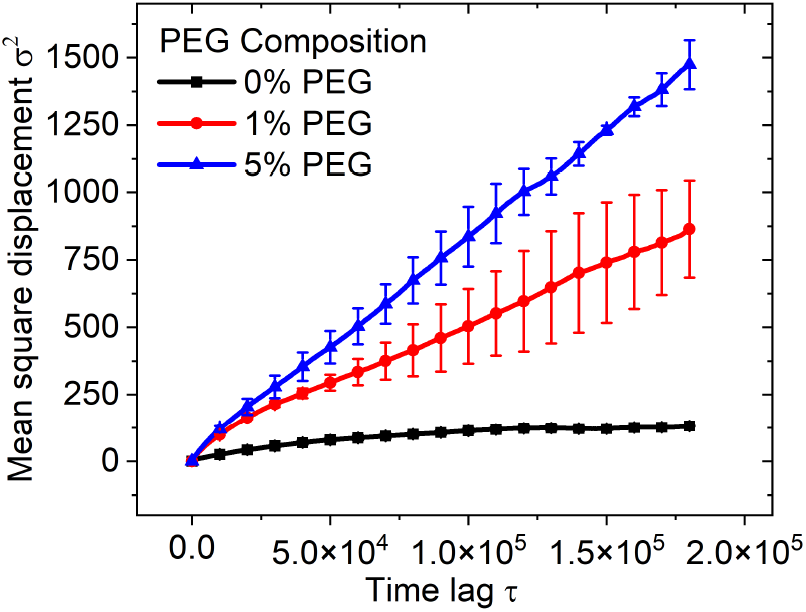
Mean square displacements of LNPs in the extracellular matrix for all PEG compositions.

We further quantify the motion of the LNPs using their diffusivities within the matrix structure. To calculate these diffusivity values, we consider the portion of the MSD curves between the time lags 5 *×* 10^4^ and 1.5 *×* 10^5^. This avoids the early timescale ballistic regime and the longer time lags with poor statistics. With this range, we calculate the slope *k* of the individual MSD curves using a linear fit. The mean diffusivities (*k/*6) of LNPs and their standard deviation are shown in Figure 8. Here, we observe that LNPs with 1% PEG show about 8 times higher mean diffusivity than the 0% PEG LNPs. Whereas the 5% PEG LNPs diffuse faster by almost 1.7 times than the LNPs with 1% PEG coatings. Here, we note that the 1% case shows larger standard deviation than the 0% or 5% cases. At 1% PEG, some LNPs achieve sufficient steric layer and start diffusing freely, while others remain adsorbed to the polymer chains, which creates a broad distribution of diffusivities. However, in other cases, a significant portion of the LNP population is either completely adsorbed or freely diffusing in the matrix, resulting in lower standard deviations. These findings suggest that the 0% PEG LNPs face strong adsorptive interactions from the HA-chained polymers, which leads to near-immobilization of LNPs in the ECM. When we introduce even 1% of PEG on the LNP surface, it generates a steric layer that is sufficient enough to break these adsorptive interactions and create a huge shift in the diffusive transport of LNPs. As we further increase this PEG composition to 5%, the multiplier in the diffusivity of LNPs is diminished. This observation indicates that at the 1% PEG case, the LNP surface is sufficiently passivated and any additional PEG molecules contribute only slightly to the steric interactions. Along with this, the effective hydrodynamic diameter of the LNP increases at 5% PEG, which offsets the diffusivity gains obtained from the reduced adhesion to the polymeric structures.

**Fig. 8.**
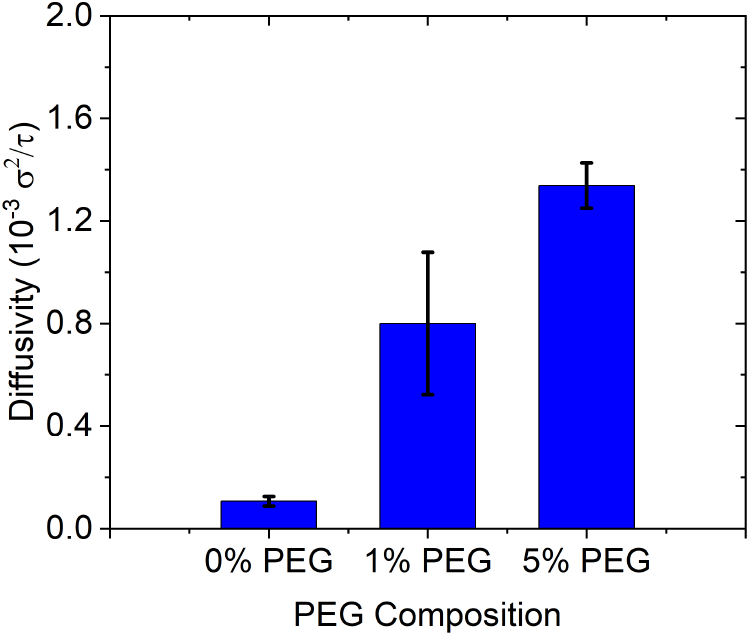
Diffusivities of LNPs in the extracellular matrix for all PEG compositions.

To investigate the change in the hydrodynamic layers on the surface of LNPs, we calculate the probability distribution as a function of distance between the polymer chain and LNP surface as shown in Figure 9. Here, we first obtain the pairwise distribution between the LNP beads and the chained polymer beads at each timestep and perform a time average of all these distributions. The profile for the 0% PEG LNPs shows a very tight packing of the LNP beads around the polymer beads. The first peak at the 2.46 nm distance immediately next to the polymer beads indicates that there is no layer of protection on the LNP surface. Here, the next peak at 3.66 nm shows secondary neighbors of the polymer beads. At 1% PEG composition, the peak locations start drifting away from the polymeric beads due to the PEG layer. Here, the primary peak (2.58 nm) starts to reduce significantly as compared to the 0% PEG, which indicates there is no tight packing of LNP beads around the polymer beads. Interestingly, the next peak at 3.78 nm shows a slightly higher probability than the primary peak. The peak value correlates with the population density at that location. A higher secondary peak indicates that the bead population is accumulating in the next shell and the region immediately next to the polymer chain is getting depleted. This structural shift is a sign of an active steric layer on the LNP surface. For the 5% PEG case, the primary peak completely vanishes, and the probability is zero till around 3 nm from the polymer chain. This profile is a diffused-out version of the 0% and 1% PEG cases. The secondary peak is entirely shifted to 4.5 nm, without any signs of packing around the polymer chains. This progressive depletion in the profiles with increasing PEG composition directly reduces the adhesive contacts between the LNP surface and polymer chains and contributes to enhanced diffusivity of the LNPs in the matrix structures.

**Fig. 9.**
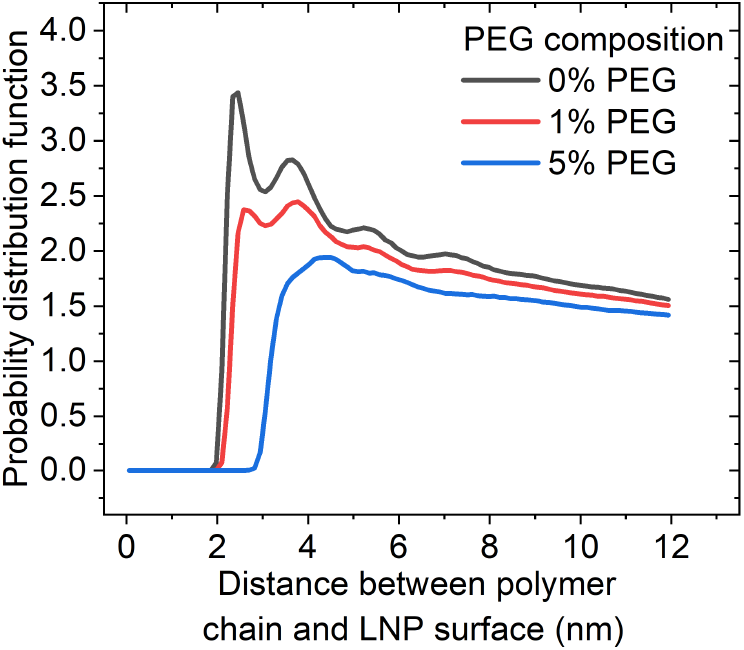
Probability distribution as a function of distance between the polymer chain and LNP surface for all PEG compositions.

Lipid nanoparticle-based therapeutics are a very promising platform for the delivery of clearance-prone molecules. Despite this, its efficacy is constrained by the adhesive interactions that the nanoparticle surface encounters during transport through the extracellular matrix. Typically particles with higher PEG compositions exhibit prolonged circulation half-life but due to these dense PEG layers they often show poor internalization in the target cells. The surface PEG density is one of the crucial design parameter in the engineering of lipid nanoparticles. The computational framework in this study, offers a way to resolve the atomistic surface interactions of LNPs with the ECM structure and quantify its impact on the diffusive transport. Our results suggest that at moderate PEG composition such as 1%, the adhesion to the matrix is disrupted and there is no excessive rise in the hydrodynamic diameter that limits diffusive transport. This framework also highlights the change in the PEG conformations from mushroom to brush-like regime where we observe the transition from depleted contact shell at 1% to complete exclusion of beads at 5% PEG. At higher PEG densities, we observe diminishing returns in the particle diffusivities as their hydrodynamic size increases. Although a denser PEG layer offers stronger steric protection, this excessive surface coverage is known to hinder the cellular uptake. The computational framework in this study offers a tool to disentangle the effect of surface PEG density from the other coupled physicochemical properties of the LNPs. This framework can be further extended to evaluate the role of other surface coatings on LNPs, lipid anchor chain length, and matrix heterogeneity to give a comprehensive predictive picture of nanoparticle transport in biological matrices.

## Conclusions

In this study, we have demonstrated the role of PEG surface density in governing these adhesive interactions and the resulting diffusive transport of LNPs through a crosslinked hyaluronic acid matrix. Our multiscale framework, which couples all-atom CpHMD atomistic interaction free energies to a highly coarse-grained model of the whole LNP, reveals that the dominant barrier to transport in the matrix is the adhesive interactions. Our analysis shows that bare LNPs pack tightly with the HA chains due to lack of any protective layer. While the introduction of even 1% PEG depletes this near-contact shell and establishes an active steric layer that disrupts these adhesive interactions. This structural change maps directly onto the diffusivity where there is eightfold increase relative to the bare LNPs. Further increase to the 5% PEG yields marginal gains of 1.7 times in the diffusivity over the LNPs with 1% PEG. This saturating behavior indicates that the LNP surface is already sufficiently passivated at low PEG densities, and that additional PEG contributes only slightly to the steric screening while increasing the effective hydrodynamic size of the LNP structure. Our in-silico multiscale framework, offers a predictive tool for accurately mapping the surface interactions of the LNPs and evaluate their impact on the diffusive transport. This computational tool, effectively complements the design of target-specific LNPs and their in-vivo experimental testing in the complex biological microenvironments.

## Data availability

The data supporting this article have been included as part of the Supplementary Information.

## Acknowledgements

This work was funded by Eli Lilly and Company. We are also grateful to the NIH and the NSF, which provided partial support for some of the computational resources through grants 1R35GM147579 and CHE-1848444/2317652.

## Notes and references

1 G. Chen, Y. Wang, R. Xie and S. Gong, Advanced drug delivery reviews, 2018, 130, 58–72.

2 W. Ho, M. Gao, F. Li, Z. Li, X.-Q. Zhang and X. Xu, Advanced healthcare materials, 2021, 10, 2001812.

3 S. Peng, B. Ouyang, Y. Men, Y. Du, Y. Cao, R. Xie, Z. Pang, S. Shen and W. Yang, Biomaterials, 2020, 231, 119680.

4 F. Zahednezhad, M. Saadat, H. Valizadeh, P. Zakeri-Milani and B. Baradaran, Journal of Controlled Release, 2019, 305, 194–209.

5 S. M. Moghimi, D. Simberg, E. Papini and Z. S. Farhangrazi, Advanced drug delivery reviews, 2020, 157, 83–95.

6 A. N. Ilinskaya and M. A. Dobrovolskaia, Toxicology and applied pharmacology, 2016, 299, 70–77.

7 D. E. Owens III and N. A. Peppas, International journal of pharmaceutics, 2006, 307, 93–102.

8 L. Tomasetti, R. Liebl, D. S. Wastl and M. Breunig, European Journal of Pharmaceutics and Biopharmaceutics, 2016, 108, 145–155.

9 D. Cahn, A. Stern, M. Buckenmeyer, M. Wolf and G. A. Duncan, ACS nano, 2024, 18, 32045–32055.

10 N. B. Shah, G. M. Vercellotti, J. G. White, A. Fegan, C. R. Wagner and J. C. Bischof, Molecular pharmaceutics, 2012, 9, 2146–2155.

11 A. E. Nel, L. Mädler, D. Velegol, T. Xia, E. M. Hoek, P. Somasundaran, F. Klaessig, V. Castranova and M. Thompson, Nature materials, 2009, 8, 543–557.

12 S.-D. Li and L. Huang, Journal of Controlled Release, 2010, 145, 178–181.

13 J. L. Perry, K. G. Reuter, M. P. Kai, K. P. Herlihy, S. W. Jones, J. C. Luft, M. Napier, J. E. Bear and J. M. DeSimone, Nano letters, 2012, 12, 5304–5310.

14 S. Xu, Z. Hu, F. Song, Y. Xu and X. Han, Molecular Therapy Methods & Clinical Development, 2025, 33, year.

15 O. Vasileva, O. Zaborova, B. Shmykov, R. Ivanov and V. Reshetnikov, Frontiers in pharmacology, 2024, 15, 1466337.

16 L. Shi, J. Zhang, M. Zhao, S. Tang, X. Cheng, W. Zhang, W. Li, X. Liu, H. Peng and Q. Wang, Nanoscale, 2021, 13, 10748–10764.

17 L. Liu, J.-H. Kim, Z. Li, M. Sun, T. Northen, J. Tang, E. Mcin-tosh, S. Karve and F. DeRosa, Nanoscale, 2025, 17, 11329–11344.

18 C. Fang, B. Shi, Y.-Y. Pei, M.-H. Hong, J. Wu and H.-Z. Chen, European Journal of Pharmaceutical Sciences, 2006, 27, 27–36.

19 V. Shalgunov, D. Zaytseva-Zotova, A. Zintchenko, T. Levada, Y. Shilov, D. Andreyev, D. Dzhumashev, E. Metelkin, A. Urusova, O. Demin et al., Journal of Controlled Release, 2017, 261, 31–42.

20 V. B. Damodaran, C. J. Fee, T. Ruckh and K. C. Popat, Langmuir, 2010, 26, 7299–7306.

21 R. Gref, M. Lück, P. Quellec, M. Marchand, E. Dellacherie, S. Harnisch, T. Blunk and R. Müller, Colloids and Surfaces B: Biointerfaces, 2000, 18, 301–313.

22 B. L. Mui, Y. K. Tam, M. Jayaraman, S. M. Ansell, X. Du, Y. Y. C. Tam, P. J. Lin, S. Chen, J. K. Narayanannair, K. G. Rajeev et al., Molecular Therapy Nucleic Acids, 2013, 2, year.

23 L. E. Waggoner, K. F. Miyasaki and E. J. Kwon, Biomaterials science, 2023, 11, 4238–4253.

24 P. de Gennes, Macromolecules, 1980, 13, 1069–1075.

25 J. McCright, C. Skeen, J. Yarmovsky and K. Maisel, Acta biomaterialia, 2022, 145, 146–158.

26 D. Makharadze, L. J. Del Valle, R. Katsarava and J. Puiggalí, International journal of molecular sciences, 2025, 26, 3102.

27 A. S. Ham, A. L. Klibanov and M. B. Lawrence, Langmuir, 2009, 25, 10038–10044.

28 P. Nakate and A. M. Ardekani, Biophysical Journal, 2026.

29 N. Aho, P. Buslaev, A. Jansen, P. Bauer, G. Groenhof and B. Hess, Journal of Chemical Theory and Computation, 2022, 18, 6148–6160.

30 A. Jansen, N. Aho, G. Groenhof, P. Buslaev and B. Hess, Journal of Chemical Information and Modeling, 2024, 64, 567–574.

31 J. A. Harris, R. Liu, V. Martins de Oliveira, E. A. Vazquez-Montelongo, J. A. Henderson and J. Shen, Journal of Chemical Theory and Computation, 2022, 18, 7510–7527.

32 V. Martins de Oliveira, Ruibin, V. Martins de Oliveira and J. Shen, Current Opinion in Structural Biology, 2022, 77, year.

33 C. A. Peeples, R. Liu and J. Shen, The Journal of Physical Chemistry B, 2024, 128, 11616–11624.

34 A. C. Thiel, M. J. Speranza, S. Jadhav, L. L. Stevens, D. K. Unruh, P. Ren, J. W. Ponder, J. Shen and M. J. Schnieders, Journal of Chemical Theory and Computation, 2024, 20, 2921–2933.

35 K. J. Colston, K. T. Faivre and S. T. Schneebeli, Nature Communications, 2025, 16, year.

36 K. J. Colston, S. C. Monsalve and S. T. Schneebeli, Molecular Pharmaceutics, 2025, 22, 7347–7358.

37 J. Philipp, A. Dabkowska, A. Reiser, K. Frank, R. Krzysz-toń, C. Brummer, B. Nickel, C. E. Blanchet, A. Sudarsan, M. Ibrahim, S. Johansson, P. Skantze, U. Skantze, S. Östman, M. Johansson, N. Henderson, K. Elvevold, B. Smedsrød, N. Schwierz, L. Lindfors and J. O. Rädler, Proceedings of the National Academy of Sciences, 2023, 120, e2310491120.

38 J. B. Simonsen and P. Larsson, Journal of Controlled Release, 2025, 384, 113879.

39 M. F. Trollmann and R. A. Böckmann, Small, 2026, 22, e11381.

40 N. B. Hamilton, S. Arns, M. Shelley, I. Bechis and J. C. Shelley, Molecular Pharmaceutics, 2024, 22, 588–593.

41 N. B. Hamilton, S. Arns, M. Shelley, I. Bechis and J. C. Shelley, Molecular Pharmaceutics, 2025, 22, 2731–2731.

42 D. J. Grzetic, N. B. Hamilton and J. C. Shelley, Molecular Pharmaceutics, 2024, 21, 4747–4753.

43 G. Settanni, W. Brill, H. Haas and F. Schmid, Macromolecular Rapid Communications, 2022, 43, 2100683.

44 J. Huang, S. Rauscher, G. Nawrocki, T. Ran, M. Feig, B. L. De Groot, H. Grubmüller and A. D. MacKerell Jr, Nature methods, 2017, 14, 71–73.

45 W. L. Jorgensen, J. Chandrasekhar, J. D. Madura, R. W. Impey and M. L. Klein, The Journal of chemical physics, 1983, 79, 926–935.

46 U. Essmann, L. Perera, M. L. Berkowitz, T. Darden, H. Lee and L. G. Pedersen, The Journal of chemical physics, 1995, 103, 8577–8593.

47 B. Hess, H. Bekker, H. J. Berendsen and J. G. Fraaije, Journal of computational chemistry, 1997, 18, 1463–1472.

48 S. Jo, T. Kim, V. G. Iyer and W. Im, Journal of computational chemistry, 2008, 29, 1859–1865.

49 W. Winter, P. Smith and S. Arnott, Journal of Molecular Biology, 1975, 99, 219–235.

50 J. S. Hub, B. L. De Groot and D. Van Der Spoel, Journal of chemical theory and computation, 2010, 6, 3713–3720.

51 H. Yuan, C. Huang, J. Li, G. Lykotrafitis and S. Zhang, Physical Review E—Statistical, Nonlinear, and Soft Matter Physics, 2010, 82, 011905.

52 A. P. Thompson, H. M. Aktulga, R. Berger, D. S. Bolintineanu, W. M. Brown, P. S. Crozier, P. J. In’t Veld, A. Kohlmeyer, S. G. Moore, T. D. Nguyen et al., Computer physics communications, 2022, 271, 108171.

