## Supplementary Information for "A multiscale modeling framework for transport of PEGylated lipid nanoparticle through the extracellular matrix"

*Non-bonded Interaction between LNP-HA beads:*

In the highly coarse-grained simulations, we keep the system at reduced temperature $k_B T = 0.23\varepsilon$, where $k_B$ is the Boltzmann's constant and $\varepsilon$ is the unit energy. We perform these computations for the physiological temperature of $310\ K$. With this value, the unit energy becomes $\varepsilon = 1.86 \times 10^{-23} kJ = 11.2\ kJ/mol$. Following our previous study [1], we keep the unit length in these simulations as $\sigma = 2\ nm$. Based on these reduced units we parametrize the 12/6 Lennard-Jones non-bonded interaction potentials. The table S1 lists these parameters in their reduced units for the different PEG compositions.

Table S1. List of interaction parameters in reduced units for the coarse-grained simulations.

| PEG Composition | Energy minimum in $\varepsilon$ units | $r_{min}$ in $\sigma$ units |
|---|---|---|
| 0% | 0.1964 | 1.211 |
| 1% | 0.1410 | 1.297 |
| 5% | 0.1391 | 1.750 |

*Non-bonded Interaction between LNP-LNP beads:*

The beads of the LNP structure interact with the anisotropic potential that depends on the distance and the relative orientation between the beads. This model is inspired from the one-particle thick membrane model developed by Yuan et al. [2]. The following set of equations describe the anisotropic form of the potential between LNP beads,

$$U(\hat{r}_{ij}, \boldsymbol{n}_i, \boldsymbol{n}_j) = \begin{cases} u_R(r) + \epsilon[1 - \phi(\hat{r}_{ij}, \boldsymbol{n}_i, \boldsymbol{n}_j)], & r < r_{\min} \\ u_{\mathrm{A}}(r)\phi(\hat{r}_{ij}, \boldsymbol{n}_i, \boldsymbol{n}_j), & r_{\min} < r < r_c \end{cases} \tag{1}$$

$$\begin{gathered} u_R(r) = \epsilon\left[\left(\frac{r_{\min}}{r}\right)^4 - 2\left(\frac{r_{\min}}{r}\right)^2\right],\ r < r_{\min}, \\ u_A(r) = -\epsilon\cos^{2\xi}\left(\frac{\pi}{2}\frac{r - r_{\min}}{r_c - r_{\min}}\right),\ r_{\min} < r < r_c, \end{gathered} \tag{2}$$

$$\begin{gathered} \phi(\hat{r}_{ij}, \boldsymbol{n}_i, \boldsymbol{n}_j) = 1 + \mu(a(\hat{r}_{ij}, \boldsymbol{n}_i, \boldsymbol{n}_j) - 1), \\ a(\hat{r}_{ij}, \boldsymbol{n}_i, \boldsymbol{n}_j) = (\boldsymbol{n}_i \times \hat{r}_{ij}) \cdot (\boldsymbol{n}_j \times \hat{r}_{ij}) + \sin\theta_0\,(\boldsymbol{n}_i - \boldsymbol{n}_j) \cdot \hat{r}_{ij} - \sin^2\theta_0\,. \end{gathered} \tag{3}$$

Here, $\phi$ function introduces the orientation dependence into the distance dependent potential $u(r)$ through the LNP bead orientation vectors $n_i$ and $n_j$. In this model we keep the bending rigidity parameter for the LNP beads constant at $\mu = 8$, for consistent mechanical properties across all the LNPs used in this study. The parameter $\sin\theta_0$ controls the curvature of the membrane surface of these LNPs, we set $\sin\theta_0 = {}^{1}/_{2r_0}$, $r_0$ is the radius of the preassembled LNP structure. Other parameters in this model are kept consistent as our previous study $\epsilon = 1\varepsilon$, $\xi = 4$, and $r_c = 2.6\sigma$ [1].

*Electrostatic interaction between beads:*

The HA chain polymers consist of negatively charged monomeric units. We model the long-range electrostatic interactions between the charged beads using Debye electrostatic potential as follows,

$$u_{\text{Debye}}(r) = \frac{Cq_iq_j}{\epsilon r} e^{-\kappa r},\ r < r_{c,\text{Debye}} \quad (4)$$

Here, $q$ is the charge on the beads and $\epsilon$ is the dielectric permittivity. $\kappa$ is the inverse of the Debye length which is kept at $1\ nm$, corresponding to the dominance of monovalent ions in the extracellular space. Here, $C$ is the energy conversion constant and $r_{c,\text{Debye}} = 5\sigma$ is the cutoff for this electrostatic potential.

*Additional All-atom CpHMD Details:*

Additional systems were simulated using different PEG lipids of varying alky lengths (C12, C14, and C18) using CpHMD, umbrella sampling, and WHAM analysis as described in the main text. There is very little difference between free energy curves when comparing different alkyl PEG lengths, as shown in Figure S1. While long longer PEG lipids are associated with slower rates of PEG lipid shedding[3], pulling the HA fibril to the surface of the model membrane along the z-axis does not model this process, resulting in the similar PMF profiles as shown below.

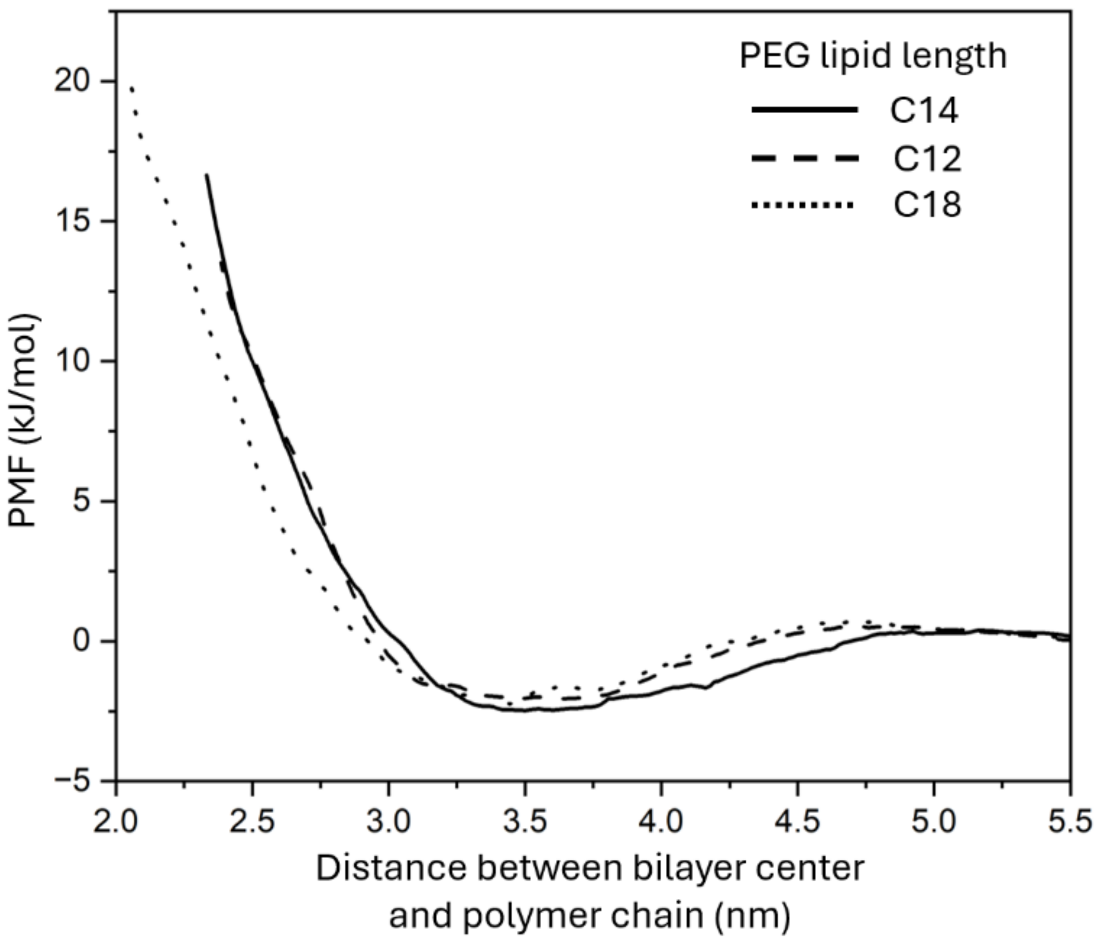


Figure S1. Superimposed free energy profiles for pulling the HA chain to the surface of a 5% PEG membrane containing different alkyl length PEG-lipids (C12 to C18). The length of the alkyl chain for the PEG lipids has minimal impact on the interaction strength between the HA chain and surface of the membrane compared to the %PEG composition at the membrane surface.

Table S1. List of components for each all-atom CpHMD simulation. All systems were roughly 5 nm x 5 nm x 12.5 nm in size.

| Component | 0% PEG (C14) | 1% PEG (C14) | 5% PEG (C12) | 5% PEG (C14) | 5% PEG (C18) |
|---|---|---|---|---|---|
| DLin-MC3-DMA | 50 | 50 | 50 | 50 | 50 |
| Cholesterol | 40 | 39 | 38 | 38 | 38 |
| DSPC | 10 | 10 | 10 | 10 | 10 |
| PEGA | 0 | 1 | 5 | 5 | 5 |
| HA subunits | 5 | 5 | 5 | 5 | 5 |
| TIP3P | 7621 | 6436 | 5997 | 6436 | 5882 |
| BUF | 500 | 500 | 500 | 500 | 500 |

References:


1. Nakate, P. and Ardekani, A.M., 2026. Modeling lipid nanoparticle transport in extracellular matrix: Effects of particle size and rigidity. Biophysical Journal, 125(4), pp.1139-1149.
2. Yuan, H., Huang, C., Li, J., Lykotrafitis, G. and Zhang, S., 2010. One-particle-thick, solvent-free, coarse-grained model for biological and biomimetic fluid membranes. Physical Review E—Statistical, Nonlinear, and Soft Matter Physics, 82(1), p.011905.
3. Ferrillo, T., Santoro, F., De Cicco, P., Guida, M., Carotenuto A., Brancaccio, D., Borrelli, F., Moore, T. L., and Quaglia, F. PEG-lipid structure controls in vitro PEG shedding, surface remodeling, and timing of lipid nanoparticle-mediated silencing. Journal of Colloid and Interface Science, **2026**, 721, 140735.
